# Graph Convolutional Neural Networks for Predicting Drug-Target Interactions

**DOI:** 10.1101/473074

**Authors:** Wen Torng, Russ B. Altman

## Abstract

Accurate determination of target-ligand interactions is crucial in the drug discovery process. In this paper, we propose a two-staged graph-convolutional (Graph-CNN) framework for predicting protein-ligand interactions. We first describe an unsupervised graph-autoencoder to learn fixed-size representations of protein pockets. Two Graph-CNNs are then trained to automatically extract features from pocket graphs and 2D molecular graphs, respectively. We demonstrate that graph-autoencoders can learn meaningful fixed-size representation for protein pockets of varying sizes and the Graph-CNN framework can effectively capture protein-ligand binding interactions without relying on target-ligand co-complexes. Across several metrics, Graph-CNNs achieved better or comparable performance to 3DCNN ligand-scoring, AutoDock Vina, RF-Score, and NNScore on common virtual screening benchmark datasets. Visualization of key pocket residues and ligand atoms contributing to the classification decisions confirms that our networks recognize meaningful interactions between pockets and ligands.

**Availability and Implementation:** Contact

**Supplementary information:**

## I. Introduction

Accurate determination of target-ligand interaction is crucial in the drug discovery process. Given a therapeutically relevant target, prioritizing the most potent compounds for the target is often the first step towards new drug development. For approved drugs, identifying unexpected off-targets can open the possibility of drug repurposing or can lead to insights for predicting and explaining observed side-effects. Currently, experimental determination of target-ligand binding affinities remains as the most accurate method for determining target-ligand interactions. However, experimental process is time-consuming and labor intensive, and cannot easily scale up to explore the diverse chemical and structural space. Computational methods have therefore been developed to perform virtual screening tasks or to predict binding affinities between target-ligand pairs.

Physic-based methods such as docking use physics-inspired rules to assess protein-ligand interactions at the atomic level^1–2^. Albeit powerful, due to their heavy dependency on atomic distances to evaluate scoring functions, these methods can be sensitive to structural error or pocket conformational changes^3^. The recent increase in protein structural data and protein-ligand interaction datasets have enabled the development of data-driven methods for target-ligand interaction predictions. These methods exploit known binding relationships and compare similarities between ligands or pockets to infer possible associations. For example, drugFEATURE^4^, evaluate pairwise pocket similarity between the query pocket and pockets with known binders and transfer binding relationship between similar targets. On the other hand, ligand-based methods evaluate molecular similarity between query compound and compounds with known targets to predict potential target interactions^5–7^. Recent methods have also integrated target-target similarity and ligand-ligand similarity to predict target-ligand binding affinities^8–9^. These methods have demonstrated promising results. However, their performance depends critically on the choice of pocket and ligand representations. Protein pockets are of varying sizes and with complex properties. It is challenging to define protein pocket similarity or to embed protein pocket as a fixed-size vector for downstream machine learning tasks without emphasizing certain aspects of pocket properties while ignoring others.

The recent success of deep learning has enabled the development of methods that can extract task-specific features directly from raw data^10^. Convolutional neural networks (CNN)^11^ are a subclass of deep learning networks that search for recurring spatial patterns in data and compose them into complex features in a hierarchical manner. Biochemical interactions start between atoms, and can be similarly aggregated over space to form complex interactions. 3DCNNs have been applied to protein structural analysis including 20 amino acid similarity analysis^12^ and protein functional site annotation^13^. Recent methods have also applied 3DCNNs to protein-ligand interaction prediction^14–15^. These methods define local boxes around pocket-ligand complexes and perform binary classification to predict whether the protein-ligand complex can interact favorably. However, since most protein-ligand pairs of interests do not have experimentally solved co-crystal structures, to generate protein-ligand co-complex for training and testing, these methods require a data-preprocessing step that artificially docks query ligands into their targets. This cross-docking step can introduce significant noises, stemming from cross-docking ligands into noncognate receptors. Furthermore, these methods are often directly trained on virtual screening benchmark datasets such as DUD-E, which has limited number and diversity of protein targets. The learned features may therefore be biased towards certain subsets of targets.

Graph convolution networks employ similar concepts of local spatial filters, but operates on graphs, and therefore has been naturally applied to 2D molecular graphs to learn small molecule representations^16–17^. In the network, each neuron is connected to its local graph neighbourhood in the previous layer through a set of learnable weights. By stacking multiple layers, each layer is looking at an increasingly larger substructure of the molecule. Simple local features can then be hierarchically composed into complex features. Kearns *et al*.^17^ applied graph convolution networks to ligand-based virtual screening and demonstrated improved performances compared to several machine learning models using Morgan fingerprints. Tsubaki et al.^18^ apply graph convolution networks and 1D-CNN to learn features from 2D molecular graphs and protein sequences, respectively.

Protein pockets consist of spatial arrangement of member residues and can go through conformational changes upon ligand binding^3^. Viewing protein pockets as graphs of residue nodes allows pockets of arbitrary size to be represented and imposes relatively weak geometry constraints. In this paper, we represent protein pocket as graphs of key residues, each taking in an input attribute that describes the local amino acid microenvironment, and propose a novel graph-convolutional framework to (1) learn meaningful protein pocket representation on a representative pocket set (2) predict protein-ligand interactions, without using protein-ligand co-complex.

Our proposed graph-convolutional framework comprises two-stages: (1) Unsupervised pocket graph-autoencoder: A graph autoencoder is first trained on a representative pocket set to embed protein pockets into fixed-size low dimensional space that perverse the input pocket properties. (2) Supervised target-ligand binding classifier: Two separate graph convolution networks are trained to learn pocket and molecular representations from pocket graphs and 2D molecular graphs, respectively. To allow the network to recognized diverse pocket features, the pocket Graph-CNN is initialized with learned weights from the pocket graph-autoencoder. The model then integrates interactions between proteins and ligands through a fully connected layer and perform binding predictions. Since features of pockets and molecules are learned through two Graph-CNN models in parallel, the model does not require protein-ligand co-complex as input. Furthermore, since the fine-tune stage training is end-to-end, driven by supervised labels, the model will automatically extract task-specific features characterizing interactions between target and molecules.

We visualized the learned pocket embedding in 2D space and show that the latent representations reflect biological pocket similarity. We then performed head-to-head comparisons of prediction performance between our Graph-CNNs, 3DCNN ligand-scoring^15^, AutoDock Vina^2^, RF-Score^19^, and NNScore^20^, on the Database of Useful Decoys: Enhanced (DUD-E)^21^ and Maximum Unbiased Validation (MUV)^22^ dataset and showed that our model achieved better or comparable performances on both datasets without requiring co-crystal structures. In addition to the conventional virtual screening setting, we further examined the ability of our model to predict binding propensity of a given compound to different targets and showed that the model can predict meaningful ligand-target binding profiles. Finally, we visualized individual contributions of each pocket residue and ligand atom to the classification decision and showed that our networks recognize meaningful interactions between protein pockets and ligands.

## II. Methods

### 2.1 Datasets

#### 2.1.1 Representative Protein Pocket Dataset

To train graph auto-encoders for extracting general features from protein pockets, we employed the druggable pocket set constructed by Liu et al.^4^ Briefly, small-molecule drugs labeled “approved” or “approved; investigational” were collected from DrugBank^23^. The drugs were mapped to their PDB^24^ IDs and high-resolution structures bound to the drug-like ligands are retrieved, resulting in 863 high-quality structures representing binding sites of 262 distinctive drugs. The above pocket set was combined with the 102 DUD-E binding pockets, resulting in a total of 965 pockets. The constructed pocket set represents a comprehensive collection of druggable binding sites.

#### 2.1.2 Database of Useful Decoys: Enhanced (DUD-E) DUD-E dataset

To enable head-to-head comparisons between Graph-CNNs, existing deep-learning-based methods and docking programs, we choose the widely-used DUD-E dataset to train our target-ligand binding classifiers. The DUD-E dataset consists of 102 targets across different protein families. On average, each target has 224 actives and over 10,000 decoys. Computational decoys are chosen such that they are physically similar but topologically dissimilar to the actives. To further improve model performance, we constructed two datasets based on the original DUD-E dataset by introducing negative pockets and experimentally validated negative ligands, as described below.

#### 2.1.2 (A) DUD-E - Pairwise interaction dataset

To train models that can predict all potential target-ligand pairwise interactions, an ideal training dataset should provide information of all positive and negative interactions between the targets and ligands in the dataset. Because the DUD-E dataset was designed for virtual screening tasks, it emphasizes on assessing binding preferences of a given target to different ligands. However, it does not provide much information regarding preferences of a given ligand to different targets. In the dataset, a given active molecule is positively associated with its true target, but not negatively associated with any targets.

To provide such information, we employ the PocketFEATURE^25^ program to assign negative binding relationships between each active ligand to a subset of the DUD-E targets. Specifically, given a protein pocket pair, the PocketFEATURE program assigns a score to quantify the extent of similarities between microenvironments in the two pockets. For a given active molecule, we calculate pairwise pocket similarity score between its true target and all the other DUD-E targets, and identify the subset of pockets in the training fold that are less similar to its true target than a predefined PocketFEATURE score threshold of −1.9 ^25^. We randomly sample 50 pockets from the identified subset and assign them as the negative pockets for the given active ligand.

#### 2.1.2 (B) DUD-E - CHEMBL assay negatives dataset

Another limitation of the original DUD-E dataset is the lack of experimentally validated negatives. Since the computational decoys are chosen to be dissimilar to actives, the benchmark excludes challenging cases where actives and negatives are similar. We construct a separate dataset that considers these challenging cases by substituting computational decoys with experimentally validated negatives from the CHEMBL^26^ database. A molecule is defined as an assay negative if its measured IC50 is higher than 50μM. To ensure the recorded interaction is direct, and only with a single protein target, only assay negatives with target confidence equal to 9 are considered. Similar to the DUDE - Pairwise interaction dataset (section 2.1.2(A)), we further added negative ligand-pocket associations using the PocketFEATURE program.

#### 2.1.3 Maximum Unbiased Validation (MUV) Dataset

To perform external validation of our models, we choose the maximum unbiased validation (MUV) data set as our independent test set. The MUV dataset consists of assay data from 17 targets, each with 30 actives and 15,000 decoys. Unlike the DUD-E dataset, the MUV dataset does not use computational decoys as negatives. Both actives and negatives are experimentally validated based on PubChem bioactivity data. Actives were selected from confirmatory screens. The decoys were selected from a primary screen for the same target. To avoid analog bias and artificial enrichment, actives were selected to be maximally spread based on simple descriptors and embedded in decoys.

To ensure the independence of our external validation dataset, we validated our models on the MUV dataset. Following procedures described in Ragoza et al.^15^, we selected MUV targets that have no more than 80% global sequence identity to any DUD-E target. Furthermore, targets with significant binding site structural similarity to any of the DUD-E target binding site were removed. Finally, only unique targets with available crystal structure of the pocket site were included. The above procedure results in eight final MUV targets. We use the same PDB structures selected by Ragoza et al.^15^ to construct our pocket graphs.

### 2.2. Input Featurization and Processing

#### Protein Pockets

We represent each protein pocket as a graph of key residues. For each of the PDB structure, we identify the pocket residues by retrieving residues that have any atom within 6 Å of the bound ligand. We define the pocket graph by viewing each of the key residue as nodes, and connecting any pair of nodes that are within 7 Å of each other with edges. For each key residue, we use the FEATURE ^27^ program to generate a fixed-size vector that describe the local amino acid microenvironment and assign the vector as the node attribute of the corresponding node. The FEAUTRE program characterizes a specified location in protein structure by dividing the local environment into six concentric shells, each of 1.25 Å in thickness. Within each shell, 80 different physicochemical properties are evaluated, resulting in a numeric vector of length 480. We normalize the FEATURE vectors such that all attributes have values between 0-1.

#### Smal1 molecules

Small molecules can be naturally represented as 2D molecular graphs, with nodes representing individual atoms and edges representing bonds. We follow the pipeline developed by Duvenaud et al.^16^ to construct the molecular graphs. Briefly, SMILES (Simplified Molecular Input Line Entry System)^28^ string encoding of each molecule is converted into a 2D molecular graph using RDKit^29^. Hydrogen atoms were treated implicitly. Each atom node is associated with simple atom descriptors including one-hot encoding of the atom’s element, the degree of the atom, the number of attached hydrogen atoms, the implicit valence, and an aromaticity indicator. The edges are associated with bond features including the bond type (single, double, triple, or aromatic), whether the bond was conjugated, and whether the bond was part of a ring.

### 2.3 Network Architecture - Graph Convolutional Neural Network

#### 2.3 (A) Unsupervised Pretraining - Pocket Graph Auto-encoder

Since the DUD-E dataset only contains 102 targets, without any unsupervised pretraining, the pocket graph convolution network will be exposed to a limited variety of protein pockets. To address this issue, we design an unsupervised framework to learn general protein pocket features on a set of 965 representative protein pockets (section 2.1.1). The unsupervised framework allows us to exploit available protein structures that have known binding sites but no sufficient binding data to learn general-purposed, fixed-sized protein pocket descriptors.

To perform unsupervised learning on graphs, we generalize basic auto-encoders^30^ onto graphs. A typical auto-encoder comprises an encoder and a decoder. The encoder project the input x into a low dimensional space (hidden layer) using the encoder weight matrix W, bias b and a non-linear activation function. The hidden layer is then transformed by the decoder weight matrix W’, bias b’ and a non-linear activation function back to its original dimension. The loss function minimizes the difference between the input signal x and the reconstructed x’. The weight matrix of the decoder may optionally be constrained by W’ = W^T^, in which case the autoencoder is said to have tied weights.

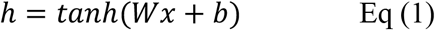

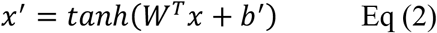

We generalize the autoencoder framework onto graphs by adapting the graph convolution operations proposed by Duvenaud et al.^16^ Specifically, our graph-convolutional autoencoder comprises two encoder-decoder stages (Figure 1), trained one after another in a greedy fashion.

**Figure 1.**
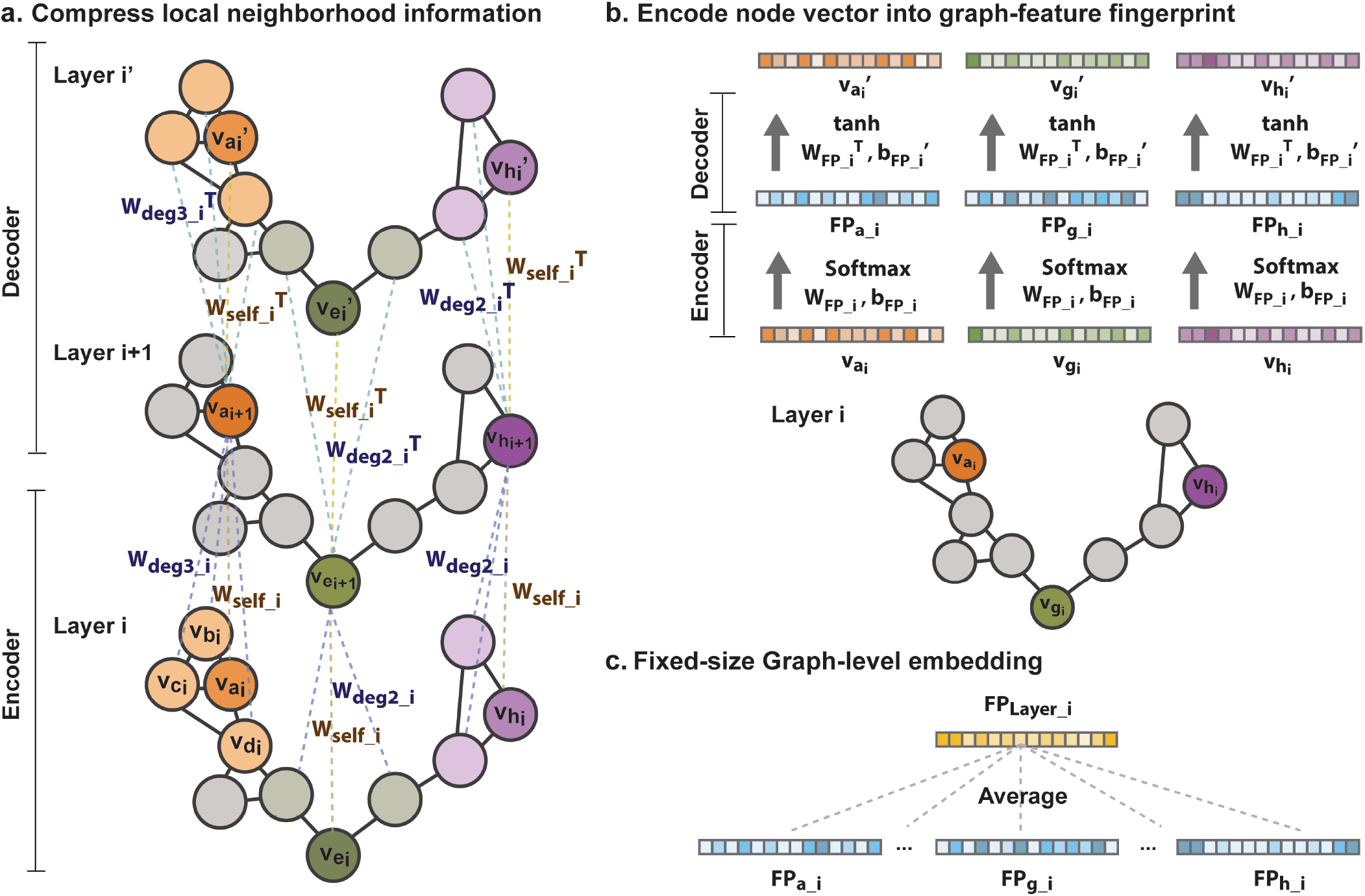
Graph convolutional autoencoder. A single graph convolutional autoencoder layer comprises two encoder-decoder stages. The two encoder-decoder stages are trained one after another in a greedy fashion. (a) The first encoder-decoder stage compresses local graph neighborhood information into low-dimensional embedding. The encoder consists of a set of graph neighborhood convolutional filters (*W_deg_y-i__*), each operates on nodes of different degrees, and a self-filter (*W_self_-_j_*), which is shared across all nodes in layer i. For each center node (*v_a_i__*)*W_deg_y-j__* convolves on the node attribute vectors of the neighborhood nodes (*v_b_i__, v_c_i__, v_d_i__*), whereas *W_self_* compresses the center node attribute vector (*v_a_i__*)). The resulting vectors are summed and transformed through ReLU to obtain *v*_*a*_*i*+1__. The decoder reconstructs the original neighborhood vectors and self-vector *v_a_i__* from *v*_*a*_*i*+1__ using the tied-weights scheme. (b) The second encoder-decoder stage compresses information from each node into a fixed-size graph-level embedding. The encoder transforms all node vectors (*v_x_i__*) in layer i to node fingerprints (*FP_x_i__*) through *W_FP_i__, b_FP_i__* and the Softmax function. The decoder reconstructs the original node vector *v_x_i__*. from the node fingerprint *FP_x_i__*, using the tied-weights scheme. (c) The resulting node fingerprints *FP_x_i__*. of all nodes in layer i are averaged to obtain a single fixed-size graph-level embedding *FP* for layer i.

#### Stage I. Compress local neighborhood information into low-dimensional embedding

The first encoder-decoder stage compresses local graph neighborhood information into low-dimensional embedding. Specifically, let *v_x_i__*. denotes the node attribute vector of node x in layer i, the first encoder and decoder stage can be described as below.

##### Encoder

The encoder consists of a set of graph neighborhood convolutional filters (*W_deg_y-i__*, *y* ∈ 0,1 … *D, where D is the maximum degree*), each operating on nodes of different degrees, and a self-filter (*W_seif_i_*), which is shared across all nodes in layer i. The encoder compresses information of graph neighborhood centered on a given node x, with degree y, by the following steps: (1) It first computes the average neighborhood vector (*v_n_i__*) of nodes in node x’s neighborhood. (2) *v_n_i__*. is then multiplied by a weight matrix *W_deg_y-i__* to obtain *z_n_i__*., where *deg_y_* specifies that the filter only operates on graph neighborhoods with degree y. (3) The node attribute vector of node x (*v_x_i__*) is multiplied by *W_self_* to obtain a transformed self-vector *z_x_i__*. (4) The two resulting vectors *z_n_i__*. and *z_x_i__*. are then summed and transformed through a ReLU^31^ function to obtain the node attribute of node x in the next layer *v*_*x*_*i*+1__.

The operations can be summarized by the below equations, where L denotes the total number of layers, H_i_ denotes the number of hidden nodes (number of filters) in graph convolution layer *i, i* ∈ 1,2.. *L*, and H_0_ represents the dimension of the input node attributes.

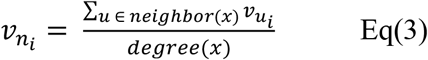

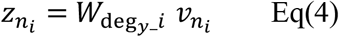

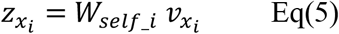

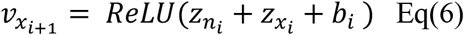

Where

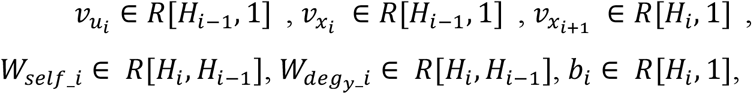

##### Decoder

The decoder reconstructs the original neighborhood vector *v_n_i__*. and self-vector *v_x_i__*. from the lower-dimension node embedding *v*_*x*_*i*+1__ using the tied-weights scheme. Specifically, to reconstruct the neighborhood vector *v_n_i__*, the decoder first multiplies *v*_*x*_*i*+1__ with 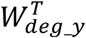 to map the hidden vector into the original dimension, adds the bias *b_n_′*, and transforms the resulting vector with a ReLU function. Similarly, to reconstruct the original node attribute *v_x_i__*, the decoder multiplies *v*_*x*_*i*+1__ with 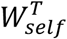, adds the bias *b_self_′* and transforms the resulting vector with a ReLU function.

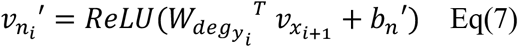

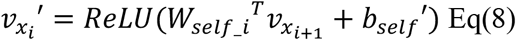

Where

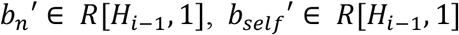

Combining the encoder and decoder, the loss function minimizes the sum of difference between *v_x_i__*. and *v_x_i__′* and between *v_n_i__* and *v_n_i__′*. Different from conventional autoencoders, the decoder needs to decode two different vectors from a single hidden vector. This process forces the hidden vector to encode information required to reconstruct both the neighborhood and self-vector information.

To conclude, the first encoder-decoder stage compresses local neighborhood information of each node in a given layer into low-dimensional embedding in the next layer. After training, the decoder layer is then discarded, and the resulting hidden node vectors are then used as input for the second encoder-decoder stage.

#### Stage II. Encode information from each node into fixed-size graph-level embedding

The second encoder-decoder stage compresses information from each node into a fixed-size graph-level embedding.

##### Encoder

All node vectors in layer i are first multiplied by the same weight matrix *W_FP_i__*, and added by bias term *b_FP_i__*. to obtain scores for each fingerprint attribute. The scores are then normalized by a Softmax^32^ function to obtain a node fingerprint (*FP_x_i__*), where the elements range from 0 to1 and sum to one. In this step, each node is classifying itself into one or multiple fingerprint categories, where each category represents a certain type of pocket property.

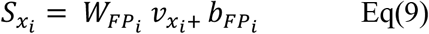

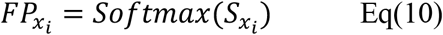

Where

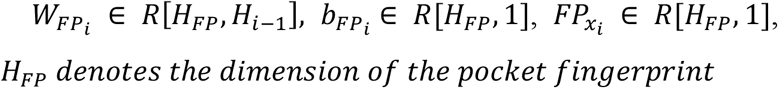

##### Decoder

The decoder reconstructs the original node vector *v_x_i__*. from the node fingerprint *FP_x_i__*, using the tied-weights scheme. Specifically, to reconstruct *v_x_i__*, the decoder multiplied *FP_x_i__*. with 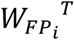, and transform the resulting vector with a tanh activation function.

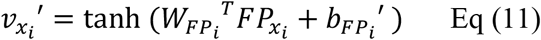

Where

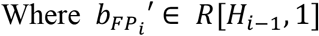

Combining the encoder and decoder, the loss function minimizes the sum of differences between *v_x__i_*. and *v_x__i_’* for all nodes in the graph. This process forces the learned node fingerprints (*FP_x_i__*) to further encode information of individual nodes (*v_x_i__*). After the training, the decoder is then discarded. The resulting node fingerprints *FP_x_i__*, of all nodes in layer i are averaged to obtain a single fixed-size graph-level embedding *FP* for layer i, regardless of the size of the graph.

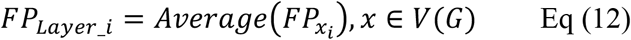

Where

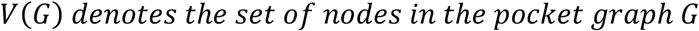

We consider the two stages as a single graph autoencoder layer. Multiple graph autoencoder layers can be stacked and trained greedily, allowing features of different complexity to be integrated through a hierarchical manner. Our final network architecture comprises two layers of graph autoencoders (Table 1). The final graph fingerprint is then obtained by summing the graph fingerprints of all layers.

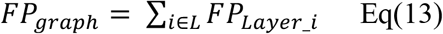

### 2.3 (B) Supervised Binding Classifier - Graph Convolution Neural Network

After training graph autoencoders to extract general-purposed protein pocket features, we proceed to construct a full model to predict drug-target interactions. Our framework comprises the following modules (Figure 2): (1) Pocket graph convolution module (2) Molecule graph convolution module (3) Interaction layer (4) Softmax classifier. Parameters of our network are summarized in Table 2.

**Figure 2.**
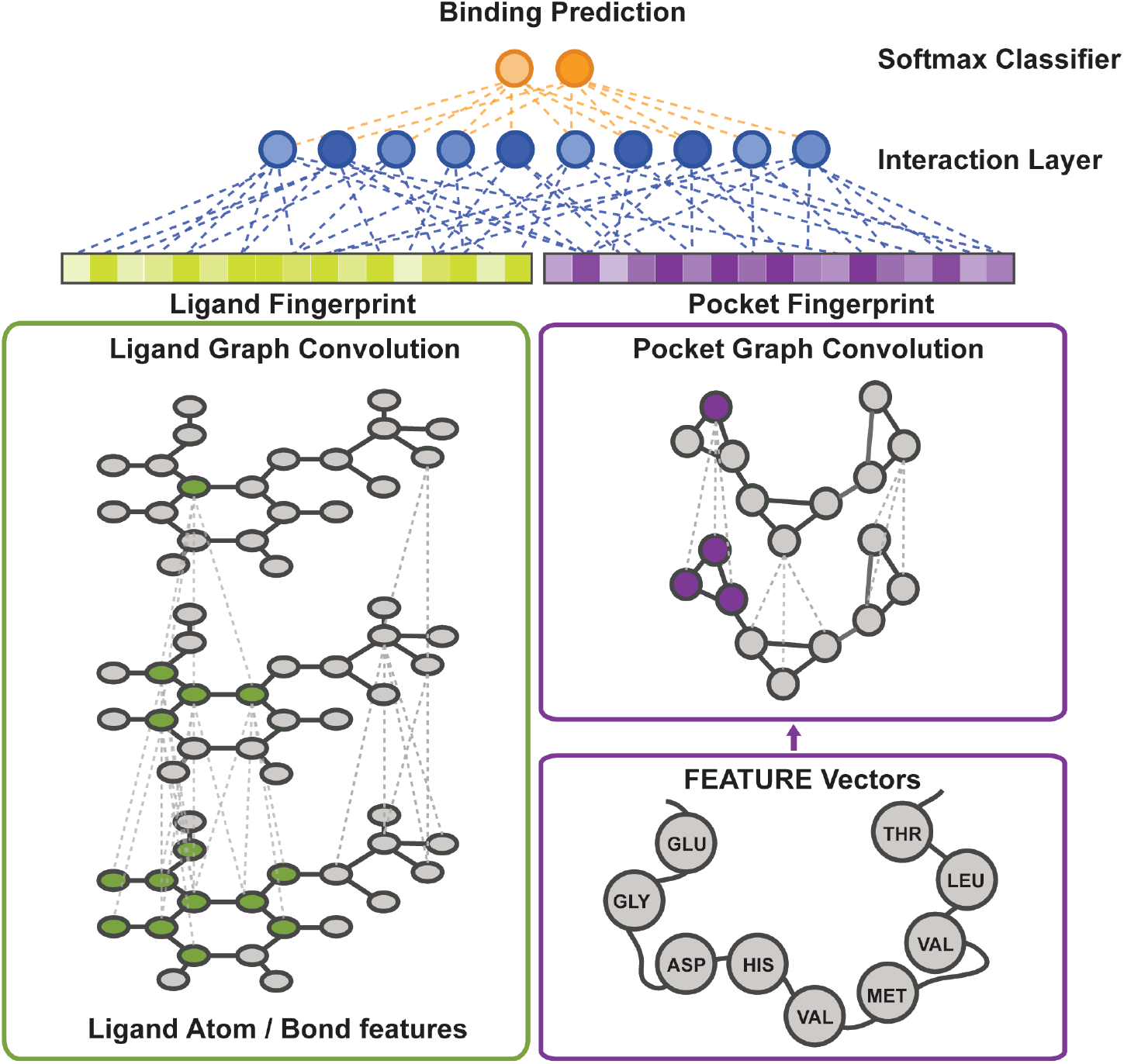
Supervised Binding Classifier. Our binding classifier comprises the following modules: (1) Pocket graph convolution module (bottom right, purple) takes in protein pockets represented as graphs, where each node corresponding to a key pocket residue and is associated with a FEATURE vector that describes the local amino acid microenvironment. The pocket graph convolution module is initialized with the learned weights from the pocket graph autoencoder and is fine-tuned to extract features from pockets that are specific to protein-ligand binding. (2) Molecule graph convolution module (bottom left, green) similarly learn features from 2D small molecular graphs. (3) The interaction layer (blue) concatenates the pocket and molecular fingerprint to generate a joint target-ligand fingerprint. The joint fingerprint is then fed to a fully-connected layer to generate the interaction hidden nodes. Each interaction hidden node represents a favorable or non-favorable interaction type between the target and ligand. (4) The Softmax classifier (orange) takes in the interaction hidden layer and calculate the class scores of “binding” or “non-binding” based on the observed favorable and non-favorable interactions.

#### (1) Pocket Graph Convolution Module

The pocket graph convolution module consists of two pocket graph convolution layers. Each pocket graph convolution layer has the same architecture as the pocket graph autoencoder (section 2.3 (A)) with the decoders removed. Specifically, the pocket graph convolution layer starts with the encoder from Stage I in the pocket graph autoencoder to extract information from local graph neighborhoods, and continues with the encoder from Stage II to compress node features into a single fixed-size graph-level embedding. This design allows us to initialize our pocket graph convolution module with the learned weights from the unsupervised graph autoencoder.

#### (2) Molecule Graph Convolution Module

We employ the graph convolution architecture developed by Duvenaud et al.^16^ to learn molecular fingerprints. Our molecular graph convolution module comprises two molecular graph convolution layers.

#### (3) Interaction Layer

The interaction layer concatenates the pocket and molecular fingerprint to generate a joint target-ligand fingerprint. A fully-connected layer takes the joint fingerprint as input and output a lower dimensional interaction hidden layer. Each node in the interaction hidden layer represents a favorable or non-favorable interaction type between the target and ligand.

#### (4) Softmax Classifier

The Softmax Classifier takes in the interaction hidden layer and calculate the class scores of “binding” or “non-binding” based on the observed favorable and non-favorable interactions.

**Table 1.**
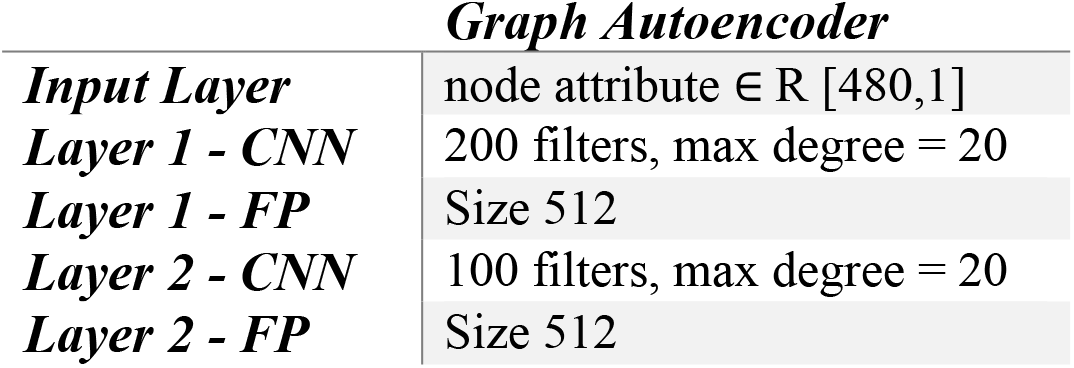
Network architecture and parameters of unsupervised pocket graph autoencoders.

| *Graph Autoencoder* |  |
| --- | --- |
| *Input Layer* | node attribute $\in \mathbb{R} [480,1]$ |
| *Layer 1 - CNN* | 200 filters, max degree = 20 |
| *Layer 1 - FP* | Size 512 |
| *Layer 2 - CNN* | 100 filters, max degree = 20 |
| *Layer 2 - FP* | Size 512 |

**Table 2.** Network architecture and parameters of target-ligand binding classifier

| *Pocket Graph-CNN* |  | *Ligand Graph-CNN* |  |
| --- | --- | --- | --- |
| *Input Layer* | node attribute $\in \mathbb{R} [480,1]$ | *Input Layer* | node attribute $\in \mathbb{R} [62,1]$<br>bond attribute $\in \mathbb{R} [6,1]$ |
| *Layer 1 - CNN* | 200 filters, max degree = 20 | *Layer 1 - CNN* | 200 filters, max degree = 6 |
| *Layer 1 - FP* | Size 512 | *Layer 1 - FP* | Size 216 |
| *Layer 2 - CNN* | 100 filters, max degree = 20 | *Layer 2 - CNN* | 100 filters, max degree = 6 |
| *Layer 2 - FP* | Size 512 | *Layer 2 - FP* | Size 216 |
| *Interaction Layer* | 728 * 100 nodes |  |  |
| *Softmax Layer* | 100*2 nodes |  |  |

### 2.4 Network Training

The unsupervised graph autoencoder was trained on the representative pocket set (section 2.1.1) in a greedy fashion until convergence. The supervised binding classifiers was trained under two different settings:

#### (1) Four-fold cross validation model - Evaluating performances on the DUDE dataset

We adopt a four-fold cross-validation strategy to train and evaluate our Graph-CNNs on the DUD-E - Pairwise interaction dataset (section 2.1.2 (A)). The folds are split between targets, where all ligands of the same target belong to the same fold. To avoid evaluating model performance on targets similar to those in the training set, we ensured that no any two folds have targets with greater than 75% global sequence identity. Within each training fold, we randomly selected three targets as our validation dataset, and choose our model hyper-parameters based on the validation performances.

The pocket graph convolution module is initialized with the pre-trained weights from section 2.3(A). The weights in the molecular graphs, the interaction layer and the final classifier are randomly initialized. During the supervised-training phase, the weights of the first pocket graph convolution layer are fixed to preserve the low-level features. We allow the higher-level weights in the pocket graph to be fine-tuned. The training process is driven by the supervised binding labels.

#### (2) Full dataset model - Evaluating performances on the independent test set: MUV dataset

To evaluate our performances on independent test dataset, we train a full model on the DUDE – CHEMBL assay negatives dataset (section 2.1.2 (B)). In addition to initializing the pocket graph convolution module with the pre-trained weights from section 2.3(A), we also pre-train a molecule graph autoencoder on all molecules in the DUDE – CHEMBL assay negatives dataset, and initialize the molecule graph convolution module with the learned weights. Similarly, weights of the first layers of the pocket and molecular graph convolution networks are fixed during the supervised training stage. The supervised model was trained for a full epoch and no rigorous attempt was made to optimize the hyper-parameters.

All models were optimized using the RMSProp algorithm^33^. The graph convolution architectures were implemented in Theano and trained on the Stanford Sherlock and Xstream servers.

### 2.5 Evaluation

#### 2.5.1 Pre-trained model: Pocket Representation

To evaluate our pre-trained pocket representation, we visualize the learned final-layer pocket fingerprints in 2D space using t-SNE^34^. We label the pockets by their corresponding SCOP^35^ families (version 2.0.7), resulting in a total of 92 classes. To facilitate better visualization, we define the majority classes as the SCOP families with more than five members, resulting in a final of 631 pockets from 38 SCOP families, and highlight each class in different colors. We additionally visualize DUDE targets and their corresponding SCOP family members, which comprises 393 pockets from 47 SCOP families.

#### 2.5.2 Supervised model: DUDE virtual screening – AUC, RE

For each target, we evaluate the virtual screening performance by the area under the receiver operating characteristic (ROC) curve metric (AUC). Specifically, binding probability of each target-ligand pair is evaluated by the corresponding test fold model. Predicted binding probabilities for all ligands for a given target is then used to calculate the AUC score for the target. For early enrichment, we employ the ROC enrichment metric (RE)^36^–^37^, chosen by Ragoza et al.^15^ Specifically, the RE score is defined as the ratio of the true positive rate (TPR) to the false positive rate (FPR) at a given FPR threshold. Here, we report the RE scores at 0.5%, 1%, 2%, and 5% FPR thresholds. We benchmark our method with 3DCNN protein-ligand scoring, Vina, and two other machine learning scoring functions, RF-Score and NNScore, using performance reported in Ragoza et al.^15^

#### 2.5.3 DUDE binding profile

Target-ligand binding interactions are known to be promiscuous. A given target can bind to multiple different ligands and vice versa. The former type of promiscuous interactions is often the focus of conventional virtual screening benchmarking datasets, whereas ligand-to-target promiscuity is often less evaluated but are nonetheless critical in predicting potential off-target effects. To assess the ability of our network to predict binding propensity of a given molecule to different targets, we use our trained networks to generate binding profiles of each active molecule in the dataset against all the DUDE targets. Specifically, we generate the binding profiles by the following steps:

(1) For a given active molecule in the DUDE dataset, we pair the active ligand with all the 102 DUDE targets, and feed the 102 ligand-target pairs into a chosen fold model. The network will then generate the “single ligand-target binding profile”, consisting of 102 binding probability scores.
(2) For a given DUDE target, we then generate the “average ligand-target binding profile” by first generating the single ligand-target binding profiles for all the actives of the given target, and average over all binding profiles.
(3) By generating the average ligand-target binding profile for all the DUDE targets and arranging them as individual rows, we can then construct a ligand-target binding propensity matrix for a given fold model. The binding propensity matrix consists of 102 rows and 102 columns, where each row correspond to the actives of the corresponding target, and each column correspond to the pocket of the corresponding target.
(4) We construct the ligand-target binding propensity matrices for all the four fold-models and evaluate the obtained matrices as follow:

#### I. Hierarchical clustering on single-fold binding propensity matrices

We perform hierarchical clustering^38^ on the rows and then the columns of each of the four binding propensity matrices to allow visualization of the groupings of targets and ligands automatically discovered by the network. The hierarchical clustering is performed using the scipy.cluster.hierarchy^39^ module.

#### II. Hierarchical clustering on test target profiles across all fold models

Since each of the four binding propensity matrices is generated using a particular fold-model, only a subset of the rows and the corresponding columns are unseen by the corresponding fold model in each fold matrix. To evaluate our model explicitly on the test cases, we generate the test-target binding propensity matrix, which is formed by the union of all the test columns from all the four binding propensity matrices. Similarly, hierarchical clustering is performed on the rows and then the columns of the matrix to discover groupings of targets and ligands.

### 2.5.4 MUV – AUC, RE

We evaluate the performances of our models on the MUV dataset using the AUC and RE metrics. In addition to comparing our performances to the 3DCNN protein-ligand scoring, Vina, RF-Score and NNScore, since ligand-based methods are known to achieve better performances than structure-based methods on the MUV dataset, we also include performances of two representative ligand-base methods extended connectivity fingerprints (ECFP4)^5^ and atom pair (AP) fingerprints^6^ for comparisons. Among them, AP is the best performing method reported in a large-scale virtual screening study of ligand-based methods conducted by Riniker et al.^7^, and ECFP4 serves as a ligand-based baseline method in our study. Performances of 3DCNN protein-ligand scoring, Vina, RF-Score and NNScore are reported as in Ragoza et al.^15^ Performances of AP and ECFP are reported as in Riniker et al.^7^

### Error analysis of MUV - Case Analysis plot

To gain further insights into the advantages and disadvantages of ligand-based and structure-based methods, we plotted the eight MUV datasets on a 2-D plot, where the X-axis represents the extent of separation of the actives from the negatives in simple chemical descriptor space and the Y-axis quantifies the average pocket similarity of the MUV target to the DUD-E pockets. Calculation of the two metrics are described in detail in Supplementary Note S1.

### Contribution from Pocket side

To quantify the extent of the pocket graph convolution module contributing to the binding predictions, for each MUV target, we replace the target protein pocket with a dummy pocket by substituting the pocket fingerprint with a zero-vector. We compare the AUC performances of all MUV targets with and without the true pockets.

### Cross-Validation on the MUV dataset

To verify whether the consistent low performance of Graph-CNNs and 3DCNN protein-ligand scoring on the MUV targets 832, 846, 852, arises from biases present in the DUD-E training dataset, we additionally trained a separate set of models using five-fold cross validation on the MUV dataset. Similar to the training of the DUD-E models, the folds are split between targets. We compare performance of the resulting models to the Graph-CNN-DUD-E models, 3DCNN protein-ligand scoring, RF-Score, NNScore, Vina, AP and ECFP4.

### 2.6 Network Visualization: Target-Ligand Interaction Map

Our target-ligand interaction map shows the contribution of pocket residues and ligand atoms to the classification decision by displaying the importance score of each pocket residue and ligand atom in heat map colors. As described in Section 2.3 (B), hidden nodes in the interaction hidden layer capture favorable or non-favorable interactions between the target and ligand. To inspect interactions captured by our network between a given input pocket and ligand pair, we identify the interaction nodes that contribute most heavily to the classification decision and visualize the contribution of each pocket residue and molecular atom to these key interactions. The identification of the key interaction nodes and the derivation of the pocket and ligand importance scores are based on stage-wise saliency map computation^42^ and are described in detail in Supplementary Note S2.

## III. Results

### 3.1 Unsupervised Pretraining - Pocket Graph Auto-encoder

To visualize protein pocket representations learned by the unsupervised graph auto-encoder, we project the learned pocket fingerprints to 2D space using t-SNE, and color the pockets by their corresponding SCOP families. Figure 3a shows the distribution of 631 pockets from 38 SCOP families that have more than five members. Figure 3b and 3c shows the distribution of the DUDE targets and the corresponding SCOP family members (401 pockets from the 48 SCOP families).

**Figure 3.**
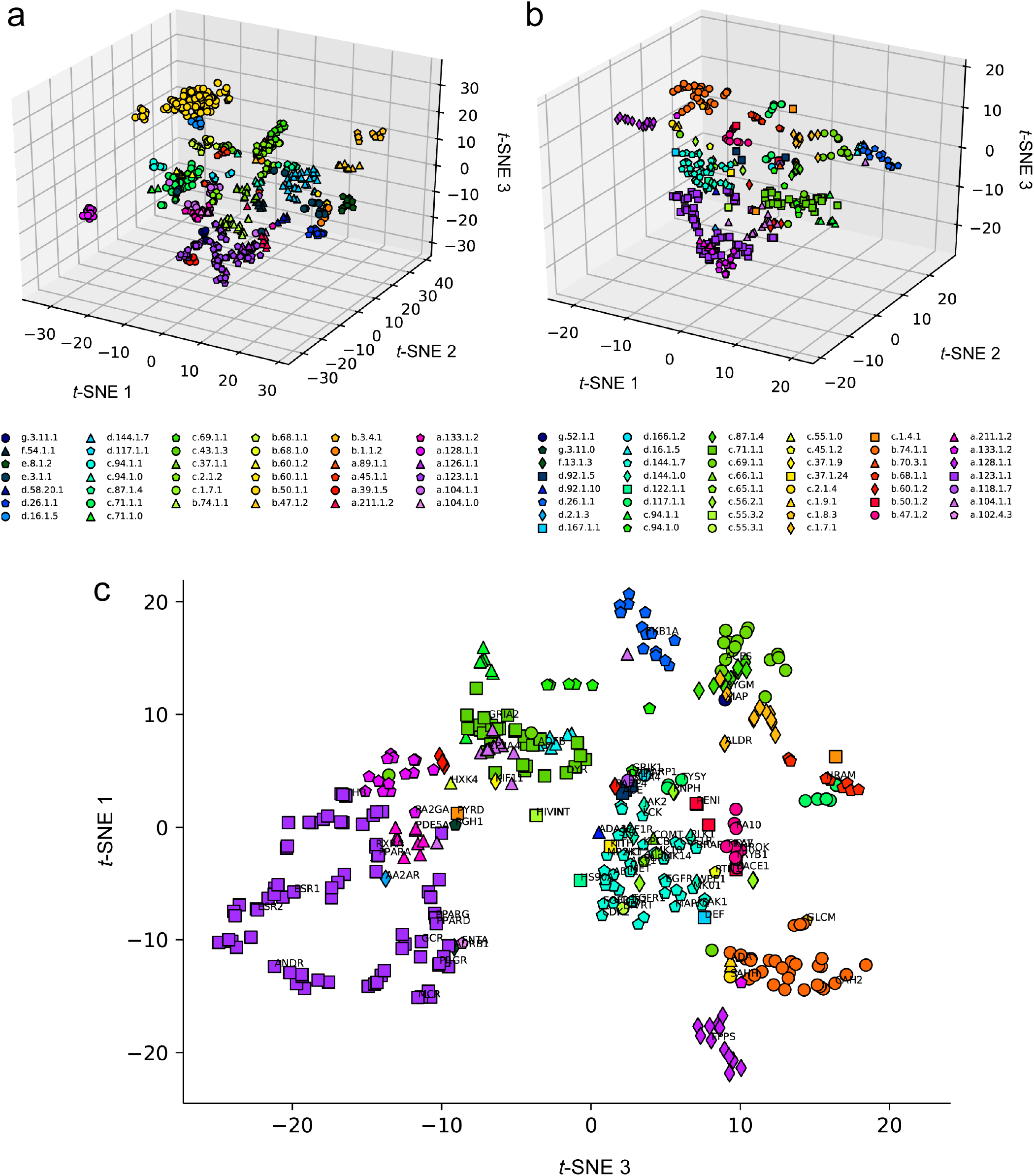
*t*-SNE visualization of the latent protein pocket space learned by the unsupervised Graph auto-encoder. **(a)** 3D *t*-SNE visualization of the distribution of 631 pockets from 38 SCOP families that have more than five members. The pockets are colored by their corresponding SCOP families. Pockets from the same SCOP family tend to lie closer to each other than to pockets in different SCOP families. **(b)** 3D *t*-SNE visualization of the distribution of the DUDE targets and the corresponding SCOP family members (401 pockets from the 48 SCOP families) in the latent pocket space. The pockets are colored by their corresponding SCOP families. **(c)** 2D projection of the 3D *t*-SNE visualization shown in (b), where the DUD-E targets are labeled with target names. The pockets are colored and marked by the same legend scheme in (b). Kinases (FAK1, BRAF, MK14, CDK2, IGF1R, CSF1R EGFR MK10, VGFR), nuclear receptors (PPARA, PPARD, PPARG, MCR GCR PRGR, ESR1, ESR2, THB), proteases (UROK, TRYB1, FA10, FA7 and TRY1) form distinct clusters in the pocket space.

### 3.2 DUDE

AUC and RE scores of Graph-CNNs, Vina, 3DCNN protein-ligand scoring, RF-Score, and NNScore on the DUD-E dataset are summarized in Table 3.

**Table 3.**
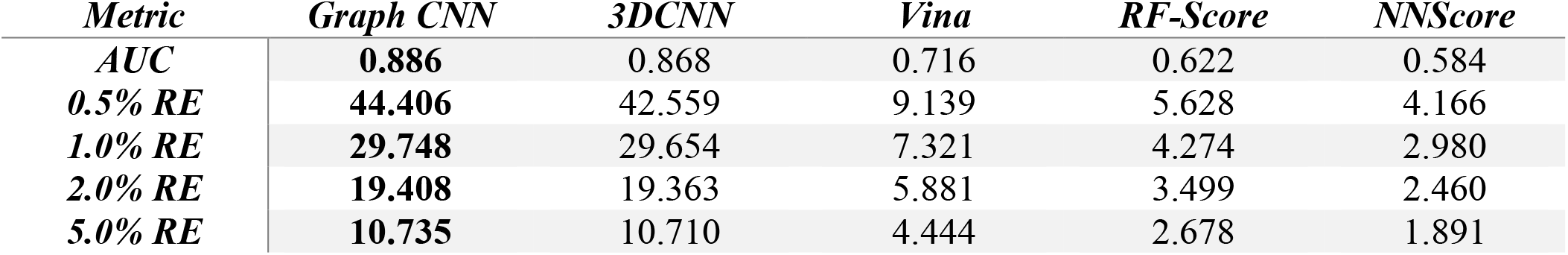
Performance of Graph-CNN, Vina, 3DCNN, RF-Score, and NNScore on the DUD-E dataset

| *Metric* | *Graph CNN* | *3DCNN* | *Vina* | *RF-Score* | *NNScore* |
| --- | --- | --- | --- | --- | --- |
| *AUC* | **0.886** | 0.868 | 0.716 | 0.622 | 0.584 |
| *0.5% RE* | **44.406** | 42.559 | 9.139 | 5.628 | 4.166 |
| *1.0% RE* | **29.748** | 29.654 | 7.321 | 4.274 | 2.980 |
| *2.0% RE* | **19.408** | 19.363 | 5.881 | 3.499 | 2.460 |
| *5.0% RE* | **10.735** | 10.710 | 4.444 | 2.678 | 1.891 |

### 3.3 DUDE Binding Profile

Figure 4 shows the binding propensity matrix of the fold-0 Graph-CNN model, where the rows correspond to the active ligands of the targets and the columns correspond to the binding pockets of the targets. Each row of the matrix contains the average ligand-target binding profile of all the actives of the corresponding target to the DUD-E targets. In other words, entry [i, j] in the matrix contains the predicted average binding propensity of active ligands of target i to target j. Note that each element in matrix are independently predicted by the model. Unlike confusion matrices, the scores of each row or each column do not sum up to 1.

**Figure 4.**
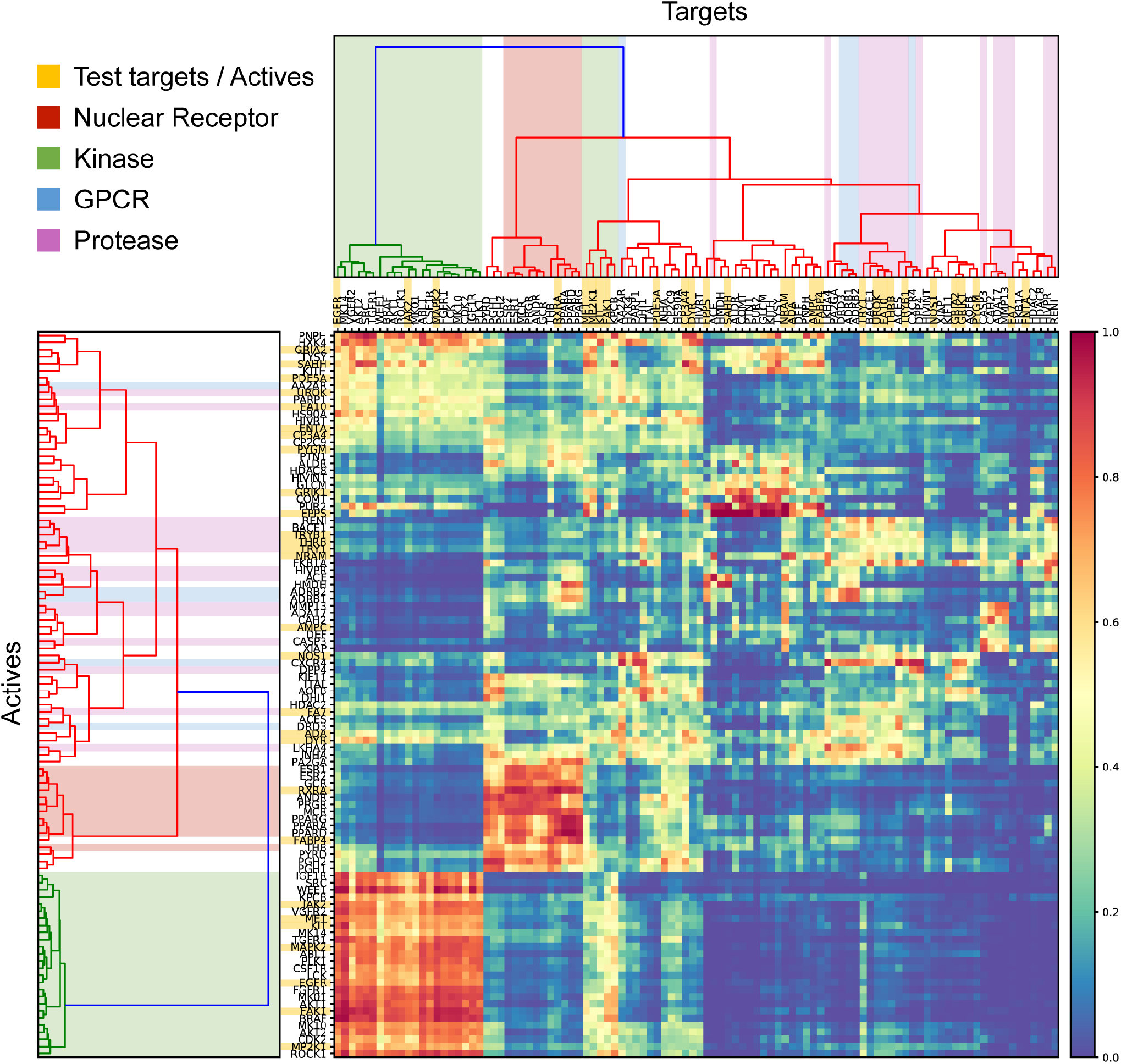
Binding propensity matrix of fold 0 model. The matrix inspects the average predicted promiscuity of ligands of a given target to all the other DUDE targets, where entry [i, j] in the matrix contains the average predicted binding propensity of active ligands of target i to target j. Hierarchical clustering on the rows reveals similarities among the ligand-target binding profiles, dividing the actives into four major clusters: (1) kinases (green), (2) nuclear receptors (red), (3) proteases (purple) and GPCRs (blue), and (4) other miscellaneous targets. Clustering on the columns similar divides the targets into four major groupings. Local high-scored blocks form at the intersections of the corresponding row and column clusters, showing that ligands are predicted to have stronger interactions with targets that are similar to its primary target compared to unrelated targets and vice versa. Test targets and ligands in fold-0 (highlighted in yellow) are clustered reasonably among the targets / ligands in the training set.

Hieratical clustering was performed on the rows and on the columns to discover similarities within the ligand-target binding profiles and within the target-ligand binding profiles. Targets of different known protein families: nuclear receptors, proteases, kinases, and GPCRs are highlighted in red, purple, green, blue respectively. Test targets and ligands in fold-0 are highlighted in yellow. Figure 5 shows the test-target binding propensity matrix across the four folds, which contains the union of the test columns from the four fold-models.

**Figure 5.**
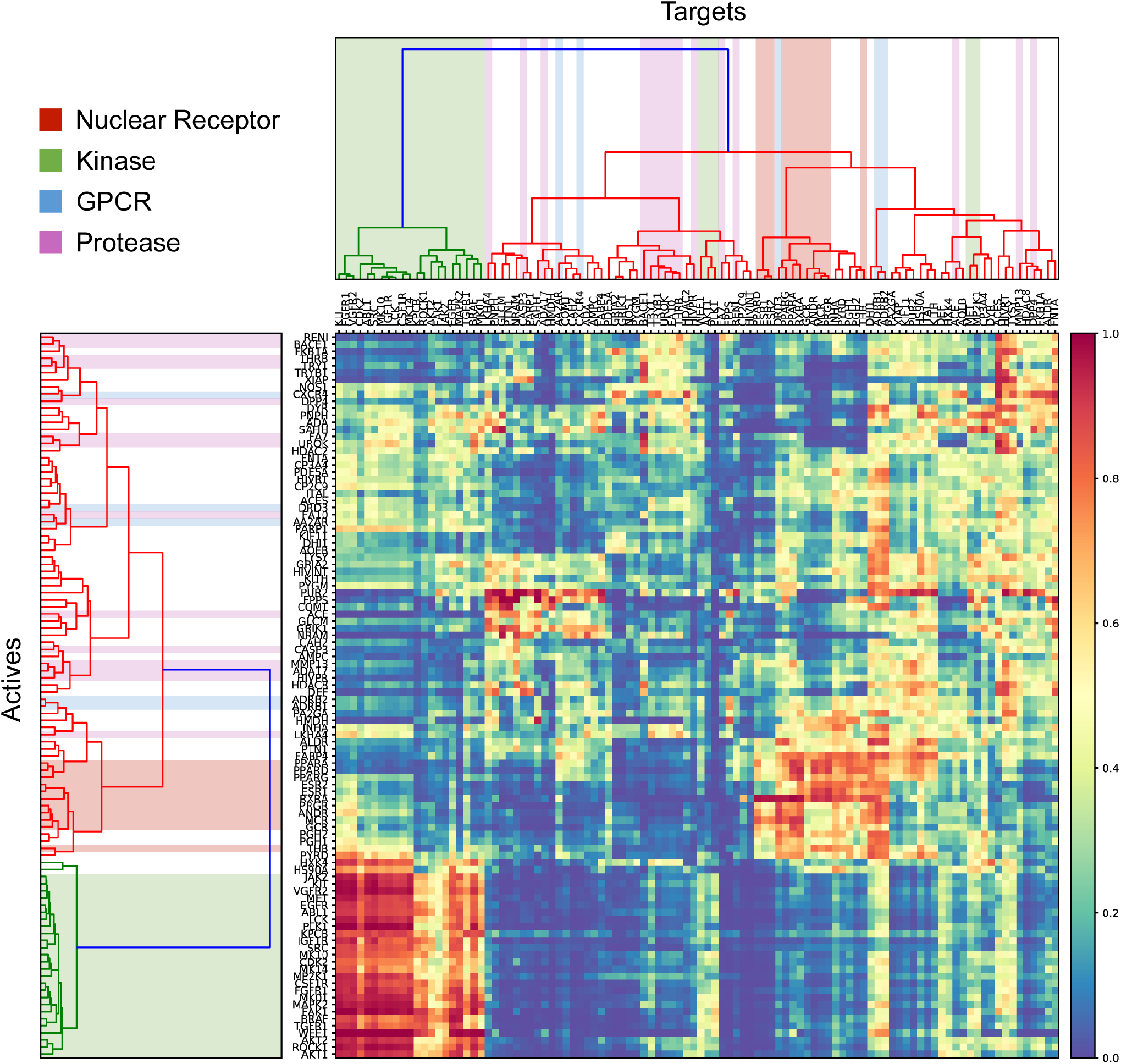
Test-target binding propensity matrix across all folds. The test-target binding propensity matrix contains the union of the test columns from the four fold-models. Although hierarchical clustering on the columns results in slightly more disperse clusters, similar targets are generally grouped together, and local high-scored blocks are observed between actives of similar targets and their corresponding targets.

### 3.4 External Validation –MUV

#### 3.4.1 Performance of Graph-CNNs on the MUV dataset

Average AUC and RE scores of Graph-CNNs, Vina, 3DCNNs, RF-Score, and NNScore on the MUV dataset are summarized in Table 4. AUC scores of each MUV target using Graph-CNNs, Vina, 3DCNNs, and ligand-based methods AP and ECFP4 are reported in Table 5.

**Table 4.** Average performance of Graph-CNN, 3DCNN, Vina, RF-Score, and NNScore on the MUV dataset

| *Metric* | *Graph CNN* | *3DCNN* | *Vina* | *RF-Score* | *NNScore* |
| --- | --- | --- | --- | --- | --- |
| *AUC* | **0.563** | 0.518 | 0.545 | 0.497 | 0.430 |
| *0.5% RE* | **3.333** | 1.667 | 0.000 | 0.000 | 0.000 |
| *1.0% RE* | **3.750** | 1.667 | 1.250 | 1.667 | 0.417 |
| *2.0% RE* | **2.500** | 1.459 | 1.459 | 1.042 | 0.417 |
| *5.0% RE* | **2.083** | 1.583 | 1.167 | 1.019 | 0.75 |

**Table 5.** Individual performance on MUV dataset using structure-based and ligand-based methods

| *Target* | *Structure-Based* |  |  | *Ligand-Based* |  |
| --- | --- | --- | --- | --- | --- |
|  | *Graph CNN* | *3DCNN* | *Vina* | *ECFP4* | *AP* |
| 859 | **0.6021** | 0.56 | 0.517 | 0.546 | 0.547 |
| 692 | **0.6234** | 0.48 | 0.413 | 0.532 | 0.588 |
| 689 | **0.7172** | 0.514 | 0.596 | 0.561 | 0.685 |
| 466 | **0.6927** | 0.663 | 0.593 | 0.494 | 0.6 |
| 548 | 0.6879 | 0.791 | 0.46 | 0.749 | **0.792** |
| 832 | 0.3859 | 0.402 | 0.61 | 0.76 | **0.771** |
| 852 | 0.4484 | 0.348 | 0.515 | 0.755 | **0.821** |
| 846 | 0.347 | 0.384 | 0.655 | 0.823 | **0.822** |

#### 3.4.2 Error analysis of MUV - Case Analysis plot

To gain further insights into the advantages and disadvantages of ligand-based and structure-based methods, we analyze properties of the ligands and pockets of the eight MUV datasets. Figure 6 shows the distribution of eight MUV target-ligand datasets according to their pocket and ligand properties, where the X-axis shows the extent of separation of the actives from the negatives of the corresponding MUV target and the Y-axis shows the average pocket similarity of the MUV target to the DUDE binding sites.

**Figure 6.**
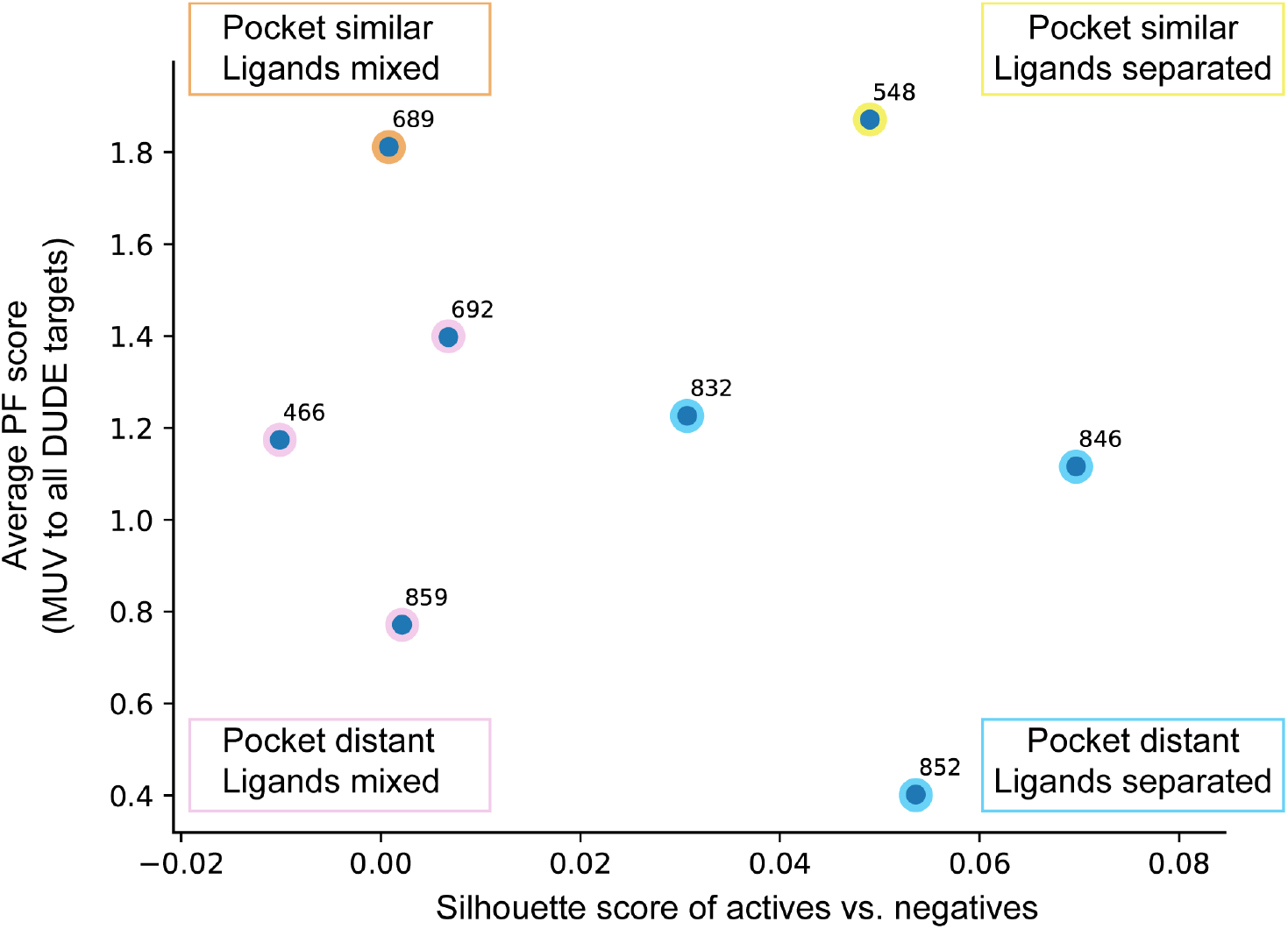
Distribution of eight MUV target-ligand datasets in the pocket-ligand property space. The X-axis shows the extent of separation of the actives from the negatives of the corresponding MUV target and the Y-axis shows the average pocket similarity of the MUV target to the DUDE binding sites. The eight MUV datasets can be divided into four categories, corresponding to the four quadrants in the ligand-pocket property space. (I) Pocket distant, Ligands mixed: actives and negatives are similar in chemical descriptor space; target pocket is distant from DUD-E targets (dataset 859,466,692) (II) Pocket similar, Ligands mixed: actives and negatives are similar in chemical descriptor space; target pocket is more similar to DUD-E targets (dataset 689) (III) Pocket distant, Ligands separated: actives and negatives are well separated in chemical descriptor space; target pocket is distant from DUD-E targets (dataset 852, 846, 832) (IV) Pocket similar, Ligands separated: actives and negatives are well separated in chemical descriptor space; target pocket is more similar to DUD-E targets (dataset 548). Ligand-based methods outperformed structure-based methods on datasets in category III: target 852, target 846, target 832. On the other hand, Graph-CNNs outperformed ligand-based methods on the hard cases in category I and II.

#### 3.4.3 Contribution of Performance from the Pocket Graphs

The AUC scores of the eight MUV datasets with dummy pockets and with true MUV target pocket are shown in Figure 7.

**Figure 7.**
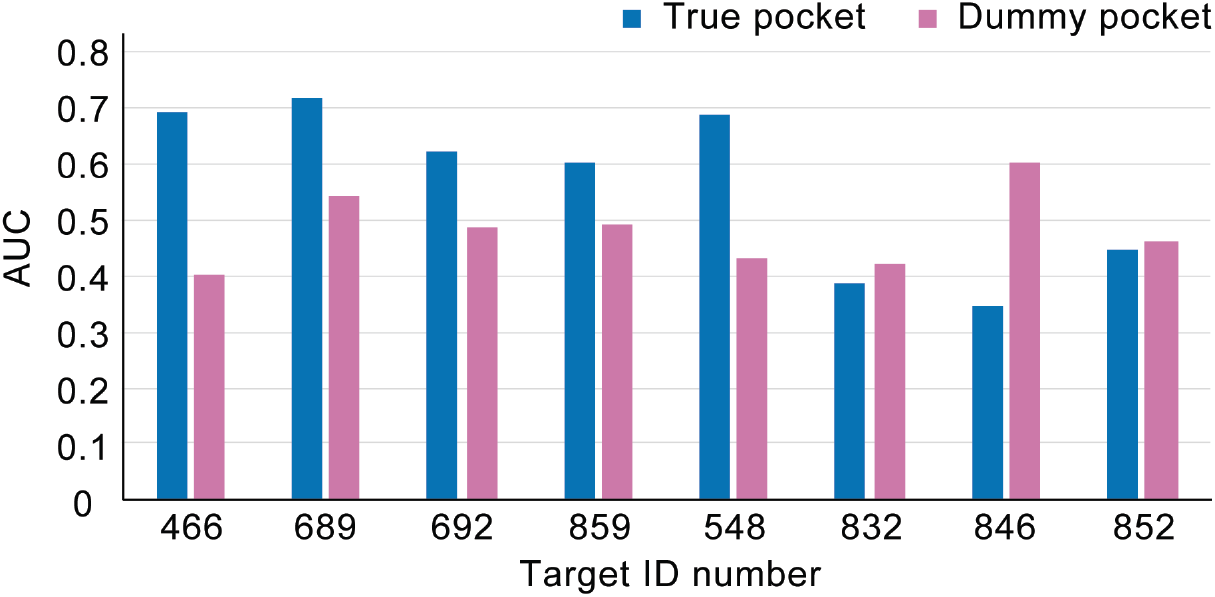
Comparisons of AUC scores on the eight MUV datasets between dummy pockets and true pockets. To examine the contribution of the pocket graph convolution module to the binding predictions, we evaluate performance of the Graph-CNN models when the true pocket is replaced with a dummy pocket. Performance of most MUV target datasets dropped substantially when the true target pockets are absent. Interestingly, target 832, 852 and 846 showed improved performance when only ligand information is provided.

#### 3.4.4 Cross-Validation on the MUV dataset

Average AUC and RE scores of the five-fold MUV-Graph-CNN models are summarized in Table 6. Comparisons of AUC scores between different methods on the MUV dataset are shown in Figure 8.

**Table 6.** Average performance of Graph-CNN-DUD-E, Graph-CNN-MUV, 3DCNN, Vina, RF-Score, and NNScore on the MUV dataset

| *Metric* | *Graph CNN-DUD-E* | *Graph CNN-MUV* | *3DCNN* | *Vina* | *RF-Score* | *NNScore* |
| --- | --- | --- | --- | --- | --- | --- |
| *AUC* | 0.563 | **0.707** | 0.518 | 0.545 | 0.497 | 0.430 |
| *0.5% RE* | 3.333 | **5.000** | 1.667 | 0.000 | 0.000 | 0.000 |
| *1.0% RE* | 3.750 | **5.833** | 1.667 | 1.250 | 1.667 | 0.417 |
| *2.0% RE* | 2.500 | **6.190** | 1.459 | 1.459 | 1.042 | 0.417 |
| *5.0% RE* | 2.083 | **4.417** | 1.583 | 1.167 | 1.019 | 0.75 |

**Figure 8.**
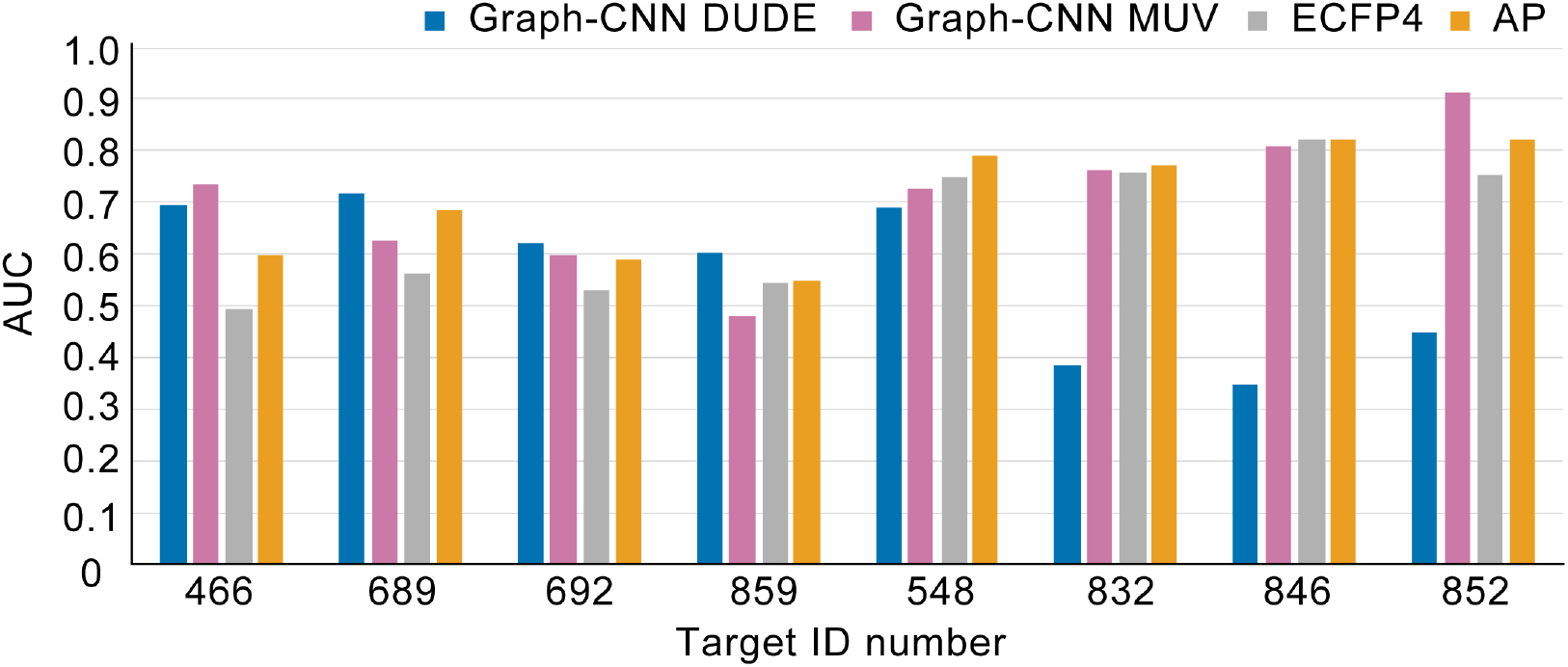
Comparisons of AUC scores between Graph-CNN-DUD-E, Graph-CNN-MUV, ECFP4, and AP on the MUV dataset. The Graph-CNN-MUV models achieve significant improvements over Graph-CNN-DUDE on targets 832, 846, and 852, achieving AUC scores comparable to AP.

### 3.5 Network Visualization

Example visualizations for target SRC, MUV-689, HDAC2, and ESR1 with their corresponding actives are shown in Figure 9. The color shows the contribution of each ligand atom and pocket residue to the classification decision. On the pocket side, the red to blue heat map spectrum highlights the most important to the least important residues. On the ligand side, the ten most important atoms are highlighted in red.

**Figure 9.**
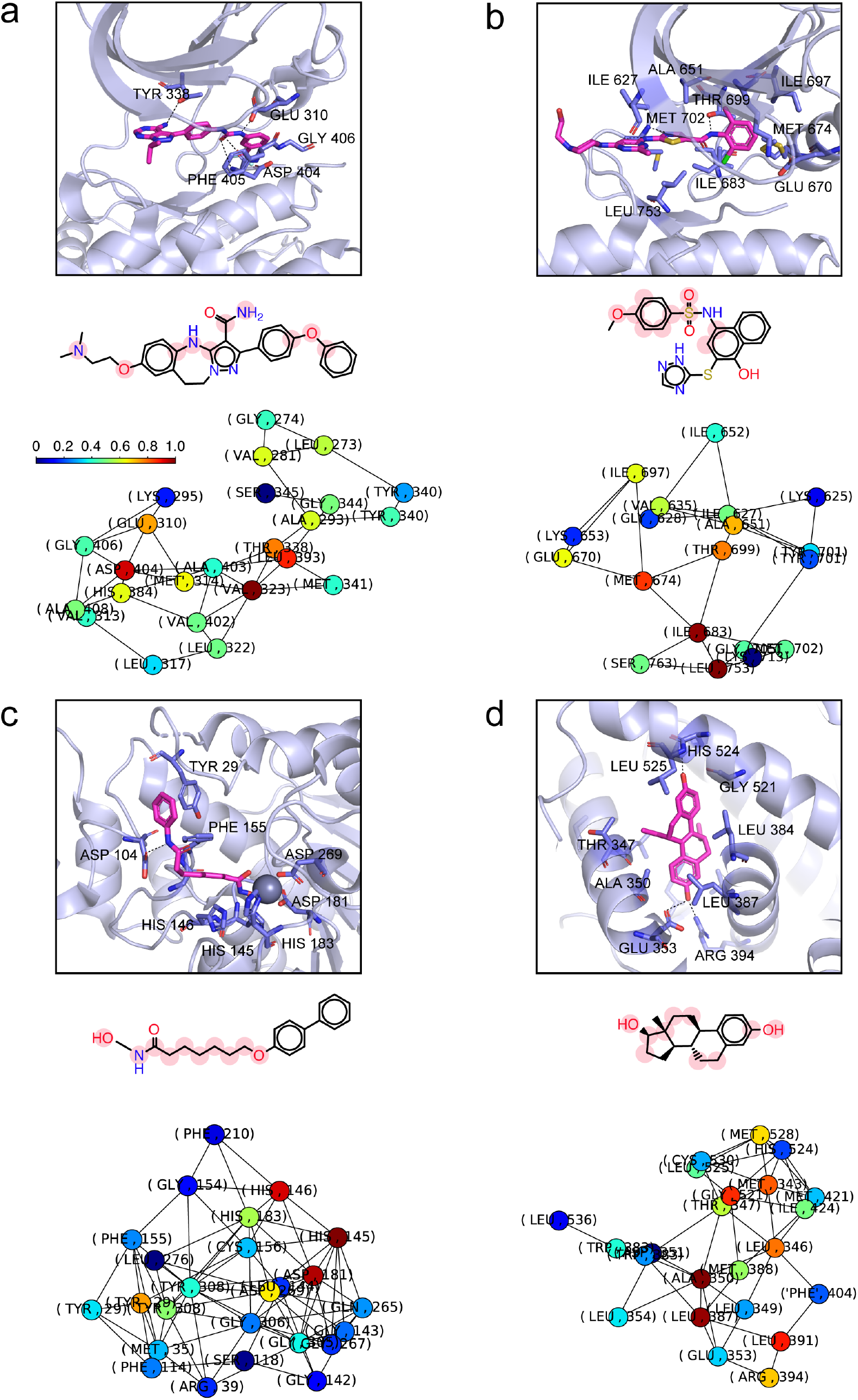
Importance visualization of pocket and ligand pairs. The color shows the contribution of each ligand atom and pocket residue to the classification decision. On the pocket side, the red to blue heat map spectrum highlights the most important to the least important residues. On the ligand side, the ten most important atoms are highlighted in red. **(a) SRC (Tyrosine-protein kinase SRC) with compound 1.** Middle and bottom: ligand and pocket importance maps for compound 1 and target SRC (PDB: 3EL8). The pocket importance map highlights residues GLU 310, TYR 338, ASP 404, VAL 323 and LEU 393. The ligand importance map highlights the nitrogen and oxygen atoms in compound 1. The identified key residues and atoms highly correspond to the observed interacting residues and ligand atoms in the co-crystalized complex of SRC and pyrazolopyrimidine 5 (Top, PDB: 3EL8). **(b) MUV – 689 (EPHA4) with compound 2.** Middle and bottom: ligand and pocket importance maps for compound 2 and EPHA4 (PDB: 2Y6O). The pocket importance map highlights residues Met 674, Thr 699, Ala 651, Ile 683, Leu 753, which highly overlap with the key pocket residues observed in the EPHA4-Dasatinib co-complex (Top, PDB: 2Y6O). The ligand interaction map highlights the benzene ring, the nearby oxygen, and the polycyclic ring in compound 2 as the key features, which may play the roles of the pyrimidine ring, the amino-thiazole ring, and the 2-chloro-6-methyl phenyl group, respectively. **(c) HDAC2 (Histone deacetylase 2) with compound 3.** Middle and bottom: ligand and pocket importance maps for compound 3 and HDAC2 (PDB: 3MAX). The pocket importance map highlights residues HIS 146, HIS 145, ASP 181, TYR 29, and ASP 269. The ligand importance map highlights the long carbon chain and the NH-OH group in compound 3. The highlighted key residues and ligand functional groups in the importance maps show high similarity to observed interactions in the HDAC2-SAHA co-complex (Top, PDB: 4LXZ). **(d) ESR1 (Estrogen receptor alpha) with compound 4.** Middle and bottom: ligand and pocket importance maps for compound 4 and ESR1 (PDB: 1SJ0). The pocket importance map highlights residues Ala 350, Leu 387, Leu 391, Leu 346, Met 343, and Gly 521, Met 528, and Arg 394. The ligand importance map highlights the two hydroxyl groups and the polycyclic ring region, which agrees with the interactions observed between ESR1 and THC (Top, PDB: 1L2I).

**Table 7.**
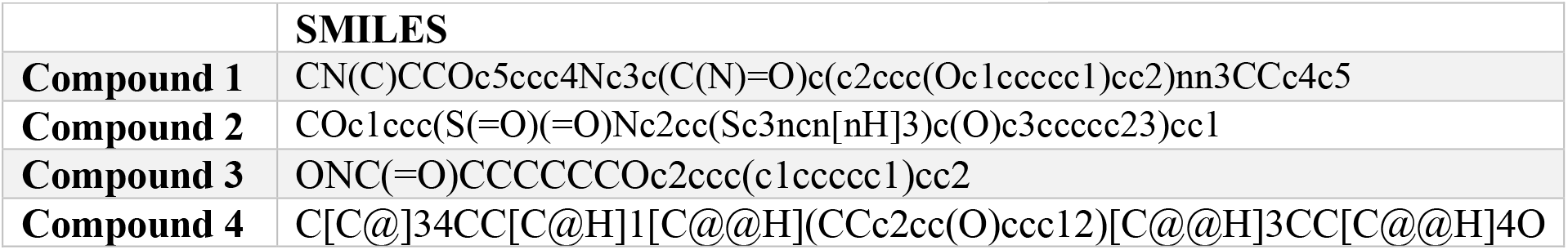
SMILES strings of compound 1 to compound 4

## IV. Discussion

### Unsupervised Pocket Representation Learning

The graph autoencoder learned meaningful pocket embeddings that reflect known pocket similarities. As shown in Figure 3a, the 631 pockets formed visible clusters, where pockets from the same SCOP family tend to lie closer to each other than to pockets in different SCOP families. Furthermore, local proximity among the clusters also reflects natural similarities between the SCOP families. For example, in the latent space, members of family b.60.1.2 (Fatty acid binding protein-like) are close to members of family b.60.1.1 (Retinol binding protein-like), both of which belonging to superfamily b.60.1 (Lipocalins).

Figure 3b and 3c shows similar trends among the DUD-E targets and their SCOP family members. For example, kinases (including FAK1, BRAF, MK14, CDK2, IGF1R, CSF1R, EGFR, MK10, VGFR) and members of the family d.144.1.7 (Protein kinases, catalytic subunit) form a visible cluster in the pocket space. Similarly, nuclear receptors (PPARA, PPARD, PPARG, MCR, GCR, PRGR, ESR1, ESR2, THB) all reside within the same region in the pocket space and are surrounded by pockets from the same SCOP family a.123.1.1 (Nuclear receptor ligand-binding domain). UROK, TRYB1, FA10, FA7 and TRY1 belong to family b.47.1.2 (Eukaryotic proteases) and form a tight cluster in the pocket space.

### DUD-E Performance and Ligand-Target Binding Profile

As shown in Table 3, the Graph-CNN models outperformed Vina, RF-score, and NNscore models and achieved comparable performance with 3DCNN protein-ligand scoring on the DUD-E dataset, without requiring co-crystal structures as input. The reported AUC and RE scores evaluate the methods on predicting the target-to-ligand binding relationships. To further assess the ability of our network to predict binding propensity of a given molecule to different targets, we constructed binding propensity matrices for each of the fold models. The matrix allows us to inspect average predicted promiscuity of ligands of a given target to all the other DUDE targets.

Figure 4 shows the binding propensity matrix of the fold-0 model. Hierarchical clustering on the rows reveals similarities among the ligand-target binding profiles, where two rows are clustered together if the corresponding ligands interact favorably with similar sets of targets. On the other hand, hierarchical clustering on the columns reveals similarities within the target-ligand binding profiles, where two columns are clustered together if the targets show strong binding propensity to similar sets of ligands. As shown in Figure 4, clustering on the rows divides the actives into four major clusters: (1) kinases (green), (2) nuclear receptors (red), (3) proteases (purple) and GPCRs (blue), and (4) other miscellaneous targets. Clustering on the columns divides the targets into similar four major groupings. Importantly, local high-scored blocks form at the intersections of the corresponding row and column clusters, showing that ligands are predicted to have stronger interactions with targets that are similar to its primary target compared to unrelated targets and vice versa. Moreover, test targets and ligands in fold-0 (highlighted in yellow) are clustered reasonably among the targets / ligands in the training set.

To further inspect the aggregated performance of the four fold-models on the test cases, we extract the test columns from all the four fold-models binding propensity matrices. Figure 5 shows the test-target binding propensity matrix, which contains the union of the test columns from the four fold-models. Although clustering on the columns results in slightly more disperse clusters, similar targets are generally grouped together, and local high-scored blocks are observed between actives of similar targets and their corresponding targets, demonstrating that the learned ligand-target binding relationship is generalizable to unseen targets.

### External Dataset Validation: MUV

We further validated our Graph-CNN models on the independent MUV benchmark. As shown in Table 4 and Table 5, performance of all methods substantially dropped on the MUV dataset. Nevertheless, the Graph-CNN models achieved the best performance among the structure-based methods across all metrics. While Vina, 3DCNN, RF-score, and NNscore have no more than two targets achieving an AUC greater than 0.6, the Graph-CNN models attain AUC greater than 0.6 for 5 out of 8 targets.

We hypothesize that the drop of performance across all methods are due to the design choices employed to construct the MUV dataset. Unlike DUD-E, the MUV dataset does not use computational decoys. Both actives and negatives are experimentally validated based on PubChem^43^ bioactivity data. Furthermore, the actives are selected to be maximally spread based on simple descriptors and embedded in decoys. Importantly, both chemical-based and cell-based assays are used to construct dataset, and thus some of the measured bioactivities may not be direct target-ligand interactions. These design choices make MUV particularly challenging for structure-based methods.

To gain insights into the advantages and disadvantages of ligand-based and structure-based methods on the MUV benchmark, we compare performance of Graph-CNN, 3DCNN, and Vina to two representative ligand-based methods AP and ECFP4 reported in a large-scale study.^7^ As shown in Table 5, Graph-CNN achieved the best performance among the 5 methods for target 859, 692, 689, 466 whereas AP achieved the best performance for target 548, 832, 852, 846.

Further analysis on the ligand and pocket properties of each MUV dataset reveals that Graph-CNNs and AP thrives at different scenarios. As shown in Figure 6, the 8 MUV datasets can be divided into four categories, corresponding to the four quadrants in the ligand-pocket property space. (I) Pocket distant, Ligands mixed: actives and negatives are similar in chemical descriptor space; target pocket is distant from DUD-E targets (dataset 859,466,692) (II) Pocket similar, Ligands mixed: actives and negatives are similar in chemical descriptor space; target pocket is more similar to DUD-E targets (dataset 689) (III) Pocket distant, Ligands separated: actives and negatives are well-separated in chemical descriptor space; target pocket is distant from DUD-E targets (dataset 852, 846, 832) (IV) Pocket similar, Ligands separated: actives and negatives are well-separated in chemical descriptor space; target pocket is more similar to DUD-E targets (dataset 548).

As expected, ligand-based methods outperformed structure-based methods on datasets in category III: target 852, target 846, target 832, where actives and negatives are well-separated in the chemical space. On the other hand, Graph-CNNs outperformed ligand-based methods on the hard cases in category I and II, where the active and negative ligands are mixed in simple molecular descriptor space, regardless of the pocket similarities between the MUV target and DUD-E targets. All methods, except Vina, performed well on category IV (target 548) where ligands are separated and the pocket similarity is relatively high.

To examine the contribution of the pocket graph convolution module to the binding predictions, we evaluate performance of the Graph-CNN models when the true pocket is replaced with a dummy pocket. As shown in Figure 7, performance of most MUV target datasets dropped substantially when the true target pockets are absent. Interestingly, target 832, 852 and 846 showed improved performance when only ligand information is provided. Two factors may have contributed to this observation: (1) These three targets had the lowest Graph-CNN performance using true pockets (2) Among the 8 MUV targets, these targets have the most separated actives and negative ligands.

The low performance on target 852, 846, and 832 are not unique to Graph-CNN models but are also observed with 3DCNN protein-ligand scoring. Graph-CNNs and 3DCNNs are both trained on the DUD-E dataset, using deep-learning-based methods. The consistent low performance of Graph-CNNs and 3DCNN protein-ligand scoring on these targets suggests that the poor performance might arise from biases in the DUD-E dataset. To investigate the hypothesis, we additionally trained Graph-CNNs on the MUV dataset using five-fold cross validation. As shown in Table 6 and Figure 8, the Graph-CNN-MUV models achieve significant improvements on the three targets, achieving AUC scores comparable to AP. This suggests that the low performance may be due to the limited coverage of interactions in the DUD-E dataset and that high-quality and large-scale datasets may be needed to improve performance of deep-learning based methods.

### Network Visualization

To visualize target-ligand interactions captured by the Graph-CNNs, we present four examples of pocket residue and ligand atom importance maps of pocket-ligand pairs, highlighting the key features contributing to the Graph-CNN classifications. Example visualizations for target SRC, MUV-689, HDAC2, and ESR1 with their corresponding active ligands are shown in Figure 9.

#### (A) SRC (Tyrosine-protein kinase SRC)

Figure 9a (middle, bottom) shows the ligand and pocket importance maps for target SRC (PDB: 3EL8^44^) and **compound 1** (Table 7). The pocket importance map shows that the positive prediction depends on residues Glu 310, Tyr 338, Asp 404, Val 323 and Leu 393. The ligand importance map shows that the decision primarily depends on the nitrogen and oxygen atoms in **compound 1**. The identified key residues and atoms highly correspond to the observed interacting residues and ligand atoms in the co-crystalized complex of SRC and pyrazolopyrimidine 5 (PDB: 3EL8.) As shown in Figure 9a (top), residue Glu 310, Tyr 338, Asp 404 form hydrogen bonds with the nitrogen and oxygen atoms in pyrazolopyrimidine 5^44^. The surrounding hydrophobic residues Val 323, Leu 393, Met 314 potentially stabilize the ligand through hydrophobic interactions. The high correspondence of the predicted and observed key residues and ligand atoms in the two target-ligand pairs suggests that the Graph-CNN is able to recognize meaningful target-ligand interactions to make predictions.

#### (B) MUV - 689 (EPHA4)

Figure 9b (middle, bottom) shows the ligand and pocket importance maps for target MUV - 689 (PDB: 2Y6O^45^) and **compound 2**. The Graph-CNN correctly classify the pocket-ligand pair as positive. The pocket importance map shows that the decision depends on residues Met 674, Thr 699, Ala 651, Ile 683, Leu 753, which highly overlap with the key pocket residues observed in the EPHA4-Dasatinib co-complex (PDB: 2Y6O). Figure 9b (top) shows the EPHA4-Dasatinib co-complex^45^. The 2- chloro-6-methyl phenyl group of dasatinib occupies the hydrophobic pocket and makes van der Waals contacts with Met 674, Ile 683, Ile 697 and Thr 699. The amino-thiazole ring has van der Waals contacts with Leu 753 and Ala 651, and also forms hydrogen bonds with Met 702. The amide group forms hydrogen bond with Thr 699. The pyrimidine ring makes contact with Ile 627. Our ligand interaction map highlights the benzene ring, the nearby oxygen, and the polycyclic ring in **compound 2** as the key features, which may play the roles of the pyrimidine ring, the amino-thiazole ring, and the 2- chloro-6- methyl phenyl group, respectively.

#### (C) HDAC2 (Histone deacetylase 2)

Figure 9c (middle, bottom) shows the ligand and pocket importance maps for target HDAC2 (PDB: 3MAX^46^) and ligand **compound 3**. The pocket importance map highlights residues His 146, His 145, Asp 181, Tyr 29, and Asp 269. The ligand importance map shows that the classification decision depends on the long carbon chain and the NH-OH group in **compound 3**. The highlighted key residues and ligand functional groups in the importance maps show high similarity to observed interactions in the HDAC2-SAHA co-complex (PDB: 4LXZ^47^). **Compound 3** belongs to the category of hydroxamic acids, which often play the role of metal chelators. Figure 9c (top) shows how SAHA, an hydroxamate inhibitor binds to the HDAC2 pocket and is coordinated to the catalytic zinc^47^. The carbon chain sits in a hydrophobic tunnel and the benzamide moiety resides at the solvent protein interface. The zinc ion is held by His 183, Asp 181, Asp 269 and chelated by the NH-OH group of the ligand. His 145, His 146, and Tyr 29 often form hydrogen bonds with hydrogens and carbonyl oxygens in the ligand^46^. The high correspondence between the predicted and observed key residues and functional groups suggests that the network was using meaningful interactions to make predictions.

#### (D) ESR1 (Estrogen receptor alpha)

Figure 9d (middle, bottom) shows the ligand and pocket importance maps for target ESR1 (PDB: 1SJ0^48^) and ligand **compound 4**. The pocket importance map shows that the Graph-CNNs made the positive prediction using residues Ala 350, Leu 387, Leu 391, Leu 346, Met 343, and Gly 521, Met 528, and Arg 394. The ligand importance map highlights the two hydroxyl groups and the polycyclic ring region, which agrees with the interactions observed between ESR1 and (R, R)-5,11-cis-diethyl-5,6,11,12-tetrahydrochrysene-2,8-diol (THC). Figure 9d (top) shows the ESR1-THC complex (PDB: 1L2I^49^), where the hydroxyl group in the phenol group forms hydrogen bonds with Arg 394 and Glu 353 and the other hydroxyl group forms hydrogen bonds with His 524 and forms van der Waals contacts with Gly 521^49^. The polycyclic ring interacts with the hydrophobic residues including Ala 350, Leu 387, Leu 384, Leu 525, Leu 391, Met 388 and Leu 346^49^.

## Conclusion

In this study, we propose a two-staged graph-convolutional framework to learn protein pocket representation and predict protein-ligand interactions. Our results show that graph-autoencoders can learn meaningful fixed-size representation for protein pockets of varying sizes that reflects protein family similarities. Such representation has the potential to enable efficient pocket similarity search, pocket classification, and can serve as input for downstream machine learning algorithms. We further demonstrate that the Graph-CNN framework can effectively capture protein-ligand binding interactions without relying on target-ligand co-complexes. Across several metrics, Graph-CNNs achieved better or comparable performance to 3DCNN ligand-scoring, AutoDock Vina, RF-Score, and NNScore on the DUD-E and MUV benchmark, and showed complementary advantages to ligand-based methods.

Our study also points out limitations in the current method and opportunities for improvement. To learn a generalizable model of protein-ligand binding, the quality of the training dataset is crucial. Virtual screening dataset often provide information about ligand binding propensity to given targets but have limited information on target preferences of given ligands. Moreover, current benchmark datasets often include limited number of targets, which only cover a subset of potential target space. Training on such dataset can result in models that perform well on prioritizing ligands for targets that are similar to those in the training set while perform badly on ranking a ligand’s true target over random targets or perform less than ideal on target-ligand interactions beyond the relationship inherent in the training data.

In this study, we impute the DUD-E with negative pocket information for each active ligand in the dataset, and initialize our pocket Graph-CNN with pretrained weights that capture prior distribution of general pocket features. We show that our model can generate meaningful ligand-target binding profiles and achieved better generalizability on the MUV dataset compared to other machine-learning and deep-learning structure-based methods. Visualization of individual contributions of each pocket residue and ligand atom also confirmed that our Graph-CNNs recognize meaningful interactions between protein pockets and ligands. However, the significant difference between performance of Graph-CNN-DUD-E and Graph-CNN-MUV models on MUV targets 832, 852 and 846 suggests that the interactions learned from the DUD-E dataset are still incomplete. We believe high quality, large-scale datasets that include both target-ligand and ligand-target binding propensities can vastly improve the performance of deep-learning-based methods on predicting ligand-target binding.

Finally, in this study, we used the FEATURE vectors to characterize microenvironments of the pocket residues. We have previously shown that 3DCNNs can extract meaningful features of amino acid microenvironments directly from raw atom distribution^12^. The proposed Graph-CNN framework can be layered on top of 3DCNNs that characterize local pocket residue environments to further enable an end-to-end framework that learns biochemical features directly from raw crystal structures. Such design can simultaneously allow detailed characterization at the local amino acid environment level and global flexibility at the pocket level.

## Acknowledgements

Computation for this study was performed on the Sherlock cluster. We would like to thank Stanford University and the Stanford Research Computing Center for providing computational resources and support that contributed to these research results. This work also used the XStream computational resource, supported by the National Science Foundation Major Research Instrumentation program (ACI-1429830).

## Funding

This work was supported by ___.

## Conflict of Interest

none declared.

